# Dog color patterns explained by modular promoters of ancient canid origin

**DOI:** 10.1101/2020.12.21.423812

**Authors:** Danika L. Bannasch, Christopher B. Kaelin, Anna Letko, Robert Loechel, Petra Hug, Vidhya Jagannathan, Jan Henkel, Petra Roosje, Marjo K. Hytönen, Hannes Lohi, Meharji Arumilli, DoGA consortium, Katie M. Minor, James R. Mickelson, Cord Drögemüller, Gregory S. Barsh, Tosso Leeb

**Affiliations:** Department of Population Health and Reproduction, School of Veterinary Medicine, University of California Davis, Davis, CA 95616 USA; Institute of Genetics, Vetsuisse Faculty, University of Bern, 3001, Bern, Switzerland; HudsonAlpha Institute for Biotechnology, Huntsville, AL, USA; Department of Genetics, Stanford University, Stanford, CA, USA; Dermfocus, University of Bern, 3001 Bern, Switzerland; VetGen, Ann Arbor, MI, 48108, USA; Division of Clinical Dermatology, Department of Clinical Veterinary Medicine, Vetsuisse Faculty, University of Bern, 3001 Bern, Switzerland; Department of Veterinary Biosciences, University of Helsinki, 00014 Helsinki, Finland; Department of Medical and Clinical Genetics, University of Helsinki, 00014 Helsinki, Finland; Folkhälsan Research Center, 00290 Helsinki, Finland; Department of Veterinary and Biomedical Sciences, University of Minnesota, Saint Paul, MN 55108, USA

## Abstract

Distinctive color patterns in dogs are an integral component of canine diversity. Color pattern differences are thought to have arisen from mutation and artificial selection during and after domestication from wolves ^1,2^ but important gaps remain in understanding how these patterns evolved and are genetically controlled ^3,4^. In other mammals, variation at the *ASIP* gene controls both the temporal and spatial distribution of yellow and black pigments ^3,5-7^. Here we identify independent regulatory modules for ventral and hair cycle *ASIP* expression, and we characterize their action and evolutionary origin. Structural variants define multiple alleles for each regulatory module and are combined in different ways to explain five distinctive dog color patterns. Phylogenetic analysis reveals that the haplotype combination for one of these patterns is shared with arctic white wolves and that its hair cycle-specific module likely originated from an extinct canid that diverged from grey wolves more than 2 million years before present. Natural selection for a lighter coat during the Pleistocene provided the genetic framework for widespread color variation in dogs and wolves.

---

A central aspect of the amazing morphologic diversity among domestic dogs are their colors and color patterns. In many mammals, specific color patterns arise through differential regulation of *Agouti* (*ASIP*), which encodes a paracrine signaling molecule that causes hair follicle melanocytes to switch from making eumelanin (black or brown pigment) to pheomelanin (yellow to nearly white pigment) ^8^. In laboratory mice, *Asip* expression is controlled by alternative promoters in specific body regions, and at specific times during hair growth, and gives rise to the light-bellied agouti phenotype, with ventral hair that is yellow and dorsal hair that contains a mixture of black and yellow pigment ^7^. Genetic variation in *ASIP* affects color pattern in many mammals; however, in dogs, the situation is still unresolved, in large part due to the complexity of different pattern types, and challenges in distinguishing whether genetic association of one or more variants represents causal variation or close linkage ^4^. Here we investigate non-coding variation in *ASIP* regulatory modules and their effect on patterning phenotypes in domestic dogs. We expand our analysis to include modern and ancient wild canids and uncover an evolutionary history in which natural selection during the Pleistocene provided a molecular substrate for color pattern diversity today.

Expression of *ASIP* promotes pheomelanin synthesis, therefore *ASIP* alleles associated with a yellow color are dominant to those associated with a black color. Although dominant yellow (DY) is common in dogs from diverse geographic locations, the most common coat pattern of modern wolves is agouti (AG) ^9^, in which the dorsum has banded hairs and the ventrum is light. Three additional color patterns are recognizable, but all have been described historically by different, inconsistent and sometimes overlapping names that predate genomic analysis (Supplementary Table 1); we refer to these as shaded yellow (SY), black saddle (BS), and black back (BB) (Fig. 1).

**Fig. 1:**
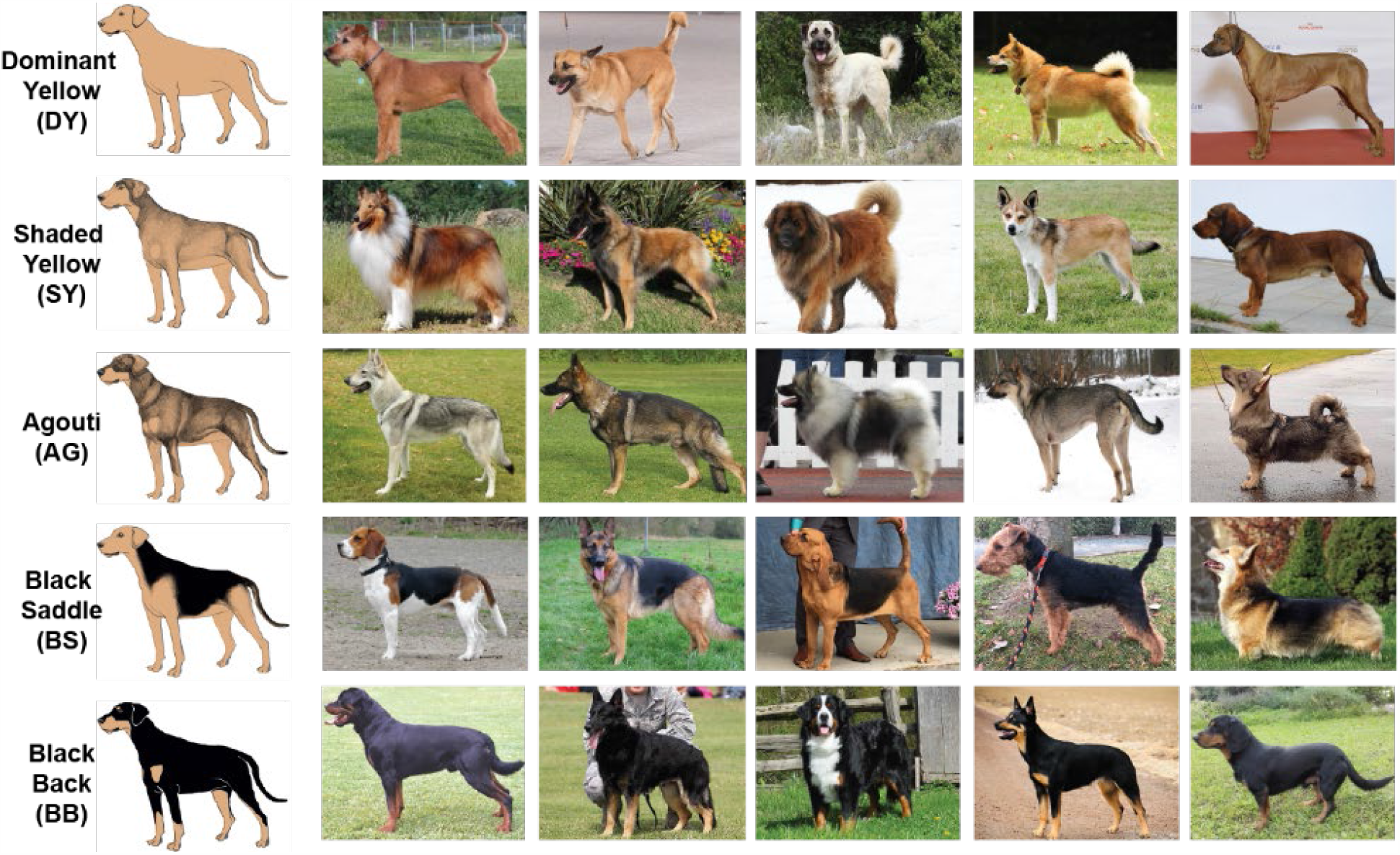
Coat patterns controlled by the *ASIP* locus. The five phenotype names proposed here are shown on the left. To the right are photographs of representative dogs of various morphological types. Length and curl of hair coat, shade of pheomelanin (red to nearly white), presence of a black facial mask and white spotting are the result of genetic variation at other loci. Patterns are displayed in order of dominance. A completely black coat caused by *ASIP* loss-of-function (recessive black) is not shown.

We collected skin RNA-seq data from dogs with different pattern phenotypes and identified three alternative untranslated first exons for dog *ASIP* (Fig. 2a, Supplementary Table 2). As described below, two of the three corresponding promoters exhibit different levels of activity and characteristic non-coding sequence variation according to dog pattern phenotype. These two promoters are orthologous to the ventral promoter (VP) and hair cycle promoter (HCP) in the laboratory mouse ^7^; however, our genetic analyses (Fig. 2) reveal that the dog VP and HCP exhibit greater regional and quantitative variation than their mouse counterparts.

**Fig. 2:**
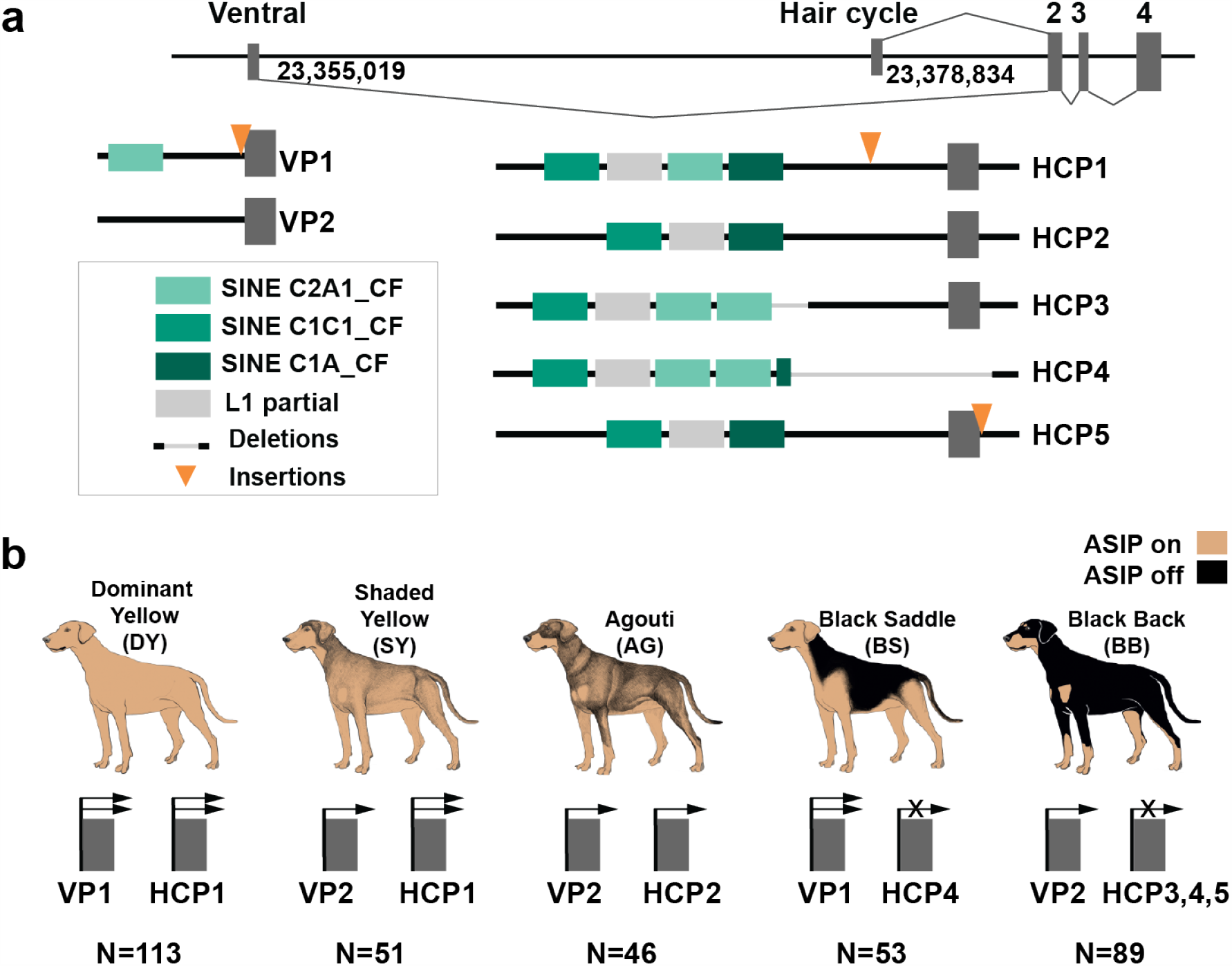
Structural variation at the *ASIP* locus in domestic dogs with different color patterns. (a) Schematic of the two relevant alternative transcription start sites and first exons (nucleotide coordinates denote their 3’-ends), together with the haplotypes observed. (b) Summary of how extended haplotype combinations are related to color pattern phenotypes. Semi-quantitative expression levels are depicted with one or two arrows or an X for no expression (Extended Data Fig. 1). N is the number of dogs for which the VP and HCP haplotype combinations accounted for *ASIP* pattern phenotype. An additional 14 dogs had a dark mask (due to an *MC1R* variant) which prevented accurate assignment of *ASIP* pattern phenotype (Extended Data Table 1, Supplementary Tables 4-7).

To better understand the relationship between promoter usage and pattern phenotypes, we inspected whole genome sequence data from 77 dog and wolf samples with known color patterns (Supplementary Table 3). We identified multiple structural variants that lie within 2 kb of the VP or HCP transcriptional start sites, confirmed their presence and identity by Sanger sequencing, and used homozygous individuals to infer two VP haplotypes and five HCP haplotypes. VP1 contains an upstream SINE element and an A-rich expansion not found in VP2 (Fig. 2a, left, Supplementary Table 1); the five HCP haplotypes differ according to the number and identity of upstream SINE elements, as well as additional insertions and deletions (Fig. 2a, right, Supplementary Table 1).

These results were extended by developing PCR-based genotyping assays for the VP and HCP structural variants, examining their association with different pattern phenotypes in 352 dogs from 34 breeds, and comparing these results to previously published variants (Extended Data Fig. 2-3, Extended Data Table 1, Supplementary Tables 4-7). As depicted in Fig. 2b, combinations of VP1 or VP2 with HCP1, 2, 3, 4, or 5 are correlated perfectly with variation in *ASIP* pattern phenotype. Because the level of *ASIP* activity is directly related to the amount of yellow pigment production, these genetic association results suggest that VP1 has greater activity than VP2, HCP1 has greater activity than HCP2, and HCP3, 4, and 5 all represent loss-of-function; indeed, the HCP4 haplotype includes a large deletion that includes the hair cycle first exon (Fig. 2a). For example, homozygotes for VP1-HCP1, VP2-HCP1, VP2-HCP2 are dominant yellow, shaded yellow and agouti, respectively (Extended Data Table 1, Supplementary Tables 4-7). Black saddle dogs are VP1-HCP4 homozygotes and most black back dogs are VP2-HCP3 homozygotes (although all three loss of function HCP haplotypes paired with VP2 can produce the black back phenotype) (Extended Data Fig.3 and Supplementary Table 7). Increased activity from the ventral promoter (VP1 vs. VP2) correlates with dorsal expansion of yellow pigment in black saddle compared to black back phenotypes (Fig. 1, 2b), which indicates that the VP and HCP haplotypes function separately from each other.

The relationship between VP and HCP variants and *ASIP* transcriptional activity was explored further using biopsies of dorsal and ventral skin (Supplementary Table 8, Extended Data Fig. 1). Read counts from RNA-seq data were consistent with expectations from the genetic association results: VP1 has greater transcriptional activity and is spatially broadened relative to VP2 (which is only expressed ventrally), HCP1 has greater transcriptional activity relative to HCP2, and no reads are detected from HCP3 or HCP4 (Fig. 2B, Extended Data Fig. 1). Taken together, these results provide a molecular explanation for *ASIP* pattern variation in dogs in which the VP and HCP haplotypes function as independent regulatory modules for their associated promoters and first exons.

Genetic relationships between variant *ASIP* regulatory modules were examined by comparing haplotypes in 18 homozygous dogs to those from 10 contemporary grey wolves (Fig. 3a, Supplementary Table 9). Overall, agouti dog haplotypes were similar to those from grey wolves. However, dominant yellow and, to a lesser extent, shaded yellow dog haplotypes were similar to those from arctic grey wolves from Ellesmere Island and Greenland, where all wolves are white (Fig. 3a, 3c). Notably, white coat color in wolves represents pale pheomelanin, as in Kermode bears or snowshoe hares ^10,11^. In the 64 kb segment that contains the VP, HCP, and coding sequence, the arctic grey wolf haplotypes are identical except for one polymorphic site, and are distinguished from dog dominant yellow haplotypes by only 6 SNVs (Extended Data Table 2). Taken together, these observations suggest a common origin of dominant yellow in dogs and white coat color in wolves without recent genetic exchange.

**Fig. 3:**
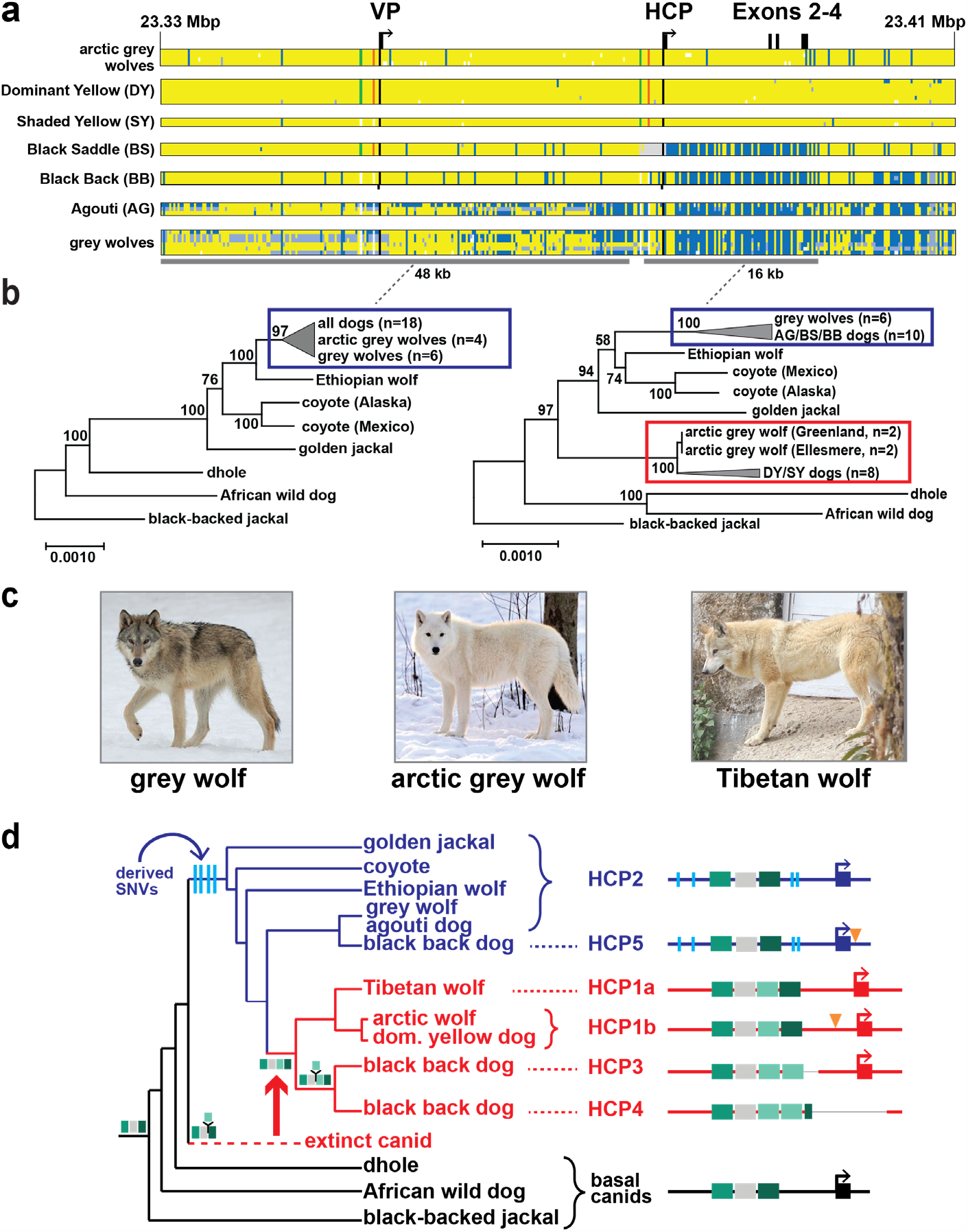
Yellow dogs and white wolves share an ancient HCP haplotype. (**a**) Genotypes at 377 SNVs (columns) at the *ASIP* locus in grey wolves and dogs (rows), coded for heterozygosity (light blue), homozygosity for the reference (yellow) or the alternate (dark blue) allele, or as missing genotypes (white). Alternate first exons (arrows) and nearby DY-associated structural variants (SINE insertions: green, polynucleotide expansions: orange) are included for reference. (**b**) Maximum likelihood phylogenies, including seven extant canid species and the dog, from 48 and 16 kb intervals upstream or downstream of the HCP, respectively. Grey wolf/dog phyletic clades are highlighted with boxes to indicate relationships that are consistent (blue) or inconsistent (red) with genome-wide phylogenies. (**c**) Images of a grey wolf, arctic grey wolf, Tibetan wolf. (**d**) A phylogeny representing distinct HCP evolutionary histories inferred from genetic variation in extant canids. Structural variants (as represented in Fig. 2) and derived SNVs (cyan) distinguish wolf-like canid (blue), ghost lineage (red), and basal canid (black) haplotypes.

The evolutionary origin of *ASIP* haplotypes was explored further by constructing maximum likelihood phylogenetic trees for dogs, wolves, and 8 additional canid species (Supplementary Table 9). Based on differences in SNV frequency, the 48 kb VP segment was considered separately from the 16 kb HCP-exon 2/3/4 segment (see supplementary text, Fig. 3a). In the VP tree, all dogs and grey wolves form a single clade, consistent with known species relationships ^12^. However, in the HCP tree, the dominant yellow and shaded yellow dogs lie in a separate clade together with arctic grey wolves; remarkably, this clade is basal to the golden jackal and distinct from other canid species (Fig. 3b, Extended Data Fig. 4, 5).

The pattern of derived allele sharing provides additional insight (Fig. 3d and Extended Data Fig. 6). As depicted in Fig. 2b and 3d, HCP2 is characterized by three small repeat elements that are shared by all canids and is therefore the ancestral form. In the branch leading to core wolf-like canids (golden jackal, coyote, Ethiopian wolf, and grey wolf), there are nine derived SNV alleles within the HCP2-exon 2/3/4 segment (Extended Data Fig. 6), four of which flank the repeat elements close to HCP2 (Fig. 3d). None of the nine derived alleles are present in the dominant yellow HCP1-exon 2/3/4 segment haplotype (which also carries an additional SINE close to HCP1; therefore this haplotype must have originated prior to the last common ancestor of golden jackals and other wolf-like canids >2 Mybp ^13^. Although the 16 kb HCP1-exon 2/3/4 segment haplotype could have originated on a branch leading to the core wolf-like canids, it would have had to persist via incomplete lineage sorting and absence of recombination for more than 2 million years and through three speciation events (supplementary text). A more likely scenario is that HCP1 represents a ghost lineage from an extinct canid (Fig. 3d, 4b) that was introduced by hybridization with grey wolves during the Pleistocene (see below), as has been suggested for an ancestor of the grey wolf and coyote ^12^, and in high altitude Tibetan and Himalayan wolves ^14^.

We expanded our analysis of VP and HCP haplotypes to a total of 45 North American and 23 Eurasian wolves, and identified a variant HCP1 haplotype in Tibetan wolves that does not extend to exon 2/3/4 and lacks the 24 bp insertion found in arctic grey wolves and dominant yellow dogs (Supplementary Table 10). The Tibetan and arctic grey wolf haplotypes are referred to as HCP1a and HCP1b, respectively (Fig. 3d, Extended Data Fig. 7, 8). The VP1-HCP1b haplotype combination is found mostly in the North American Arctic in a distribution parallel to that of white coat color (Extended Data Fig. 7a) ^15^. This haplotype combination is not observed in Eurasia, although one similar to shaded yellow, VP2-HCP1a, was observed in seven light-colored wolves from Tibet or Inner Mongolia (Fig. 3d, Extended Data Fig.7b) ^16^.

Additional insight into the demographic history of these haplotypes emerges from analysis of ancient dog (n=5) and grey wolf (n=2) WGS data, dated 4,000 – 35,000 ybp (Supplementary text and Supplementary Table 10), in which both forms of the VP (VP1 and VP2), and four forms of the HCP (HCP1a, HCP1b, HCP2, HCP4) were observed in various combinations (Fig. 4a, Extended Data Fig. 8). Ancient wolves from the Lake Taimyr and Yana River areas of Arctic Siberia had at least one HCP1 haplotype, while ancient dogs from central Europe, Ireland, and Siberia carried HCP1a, HCP1b, and HCP4, respectively (Supplementary Table 10). Thus, diversity in *ASIP* regulatory sequences responsible for color variation today was apparent by 35,000 ybp in ancient wolves and by 9,500 ybp in ancient dogs.

**Fig. 4:**
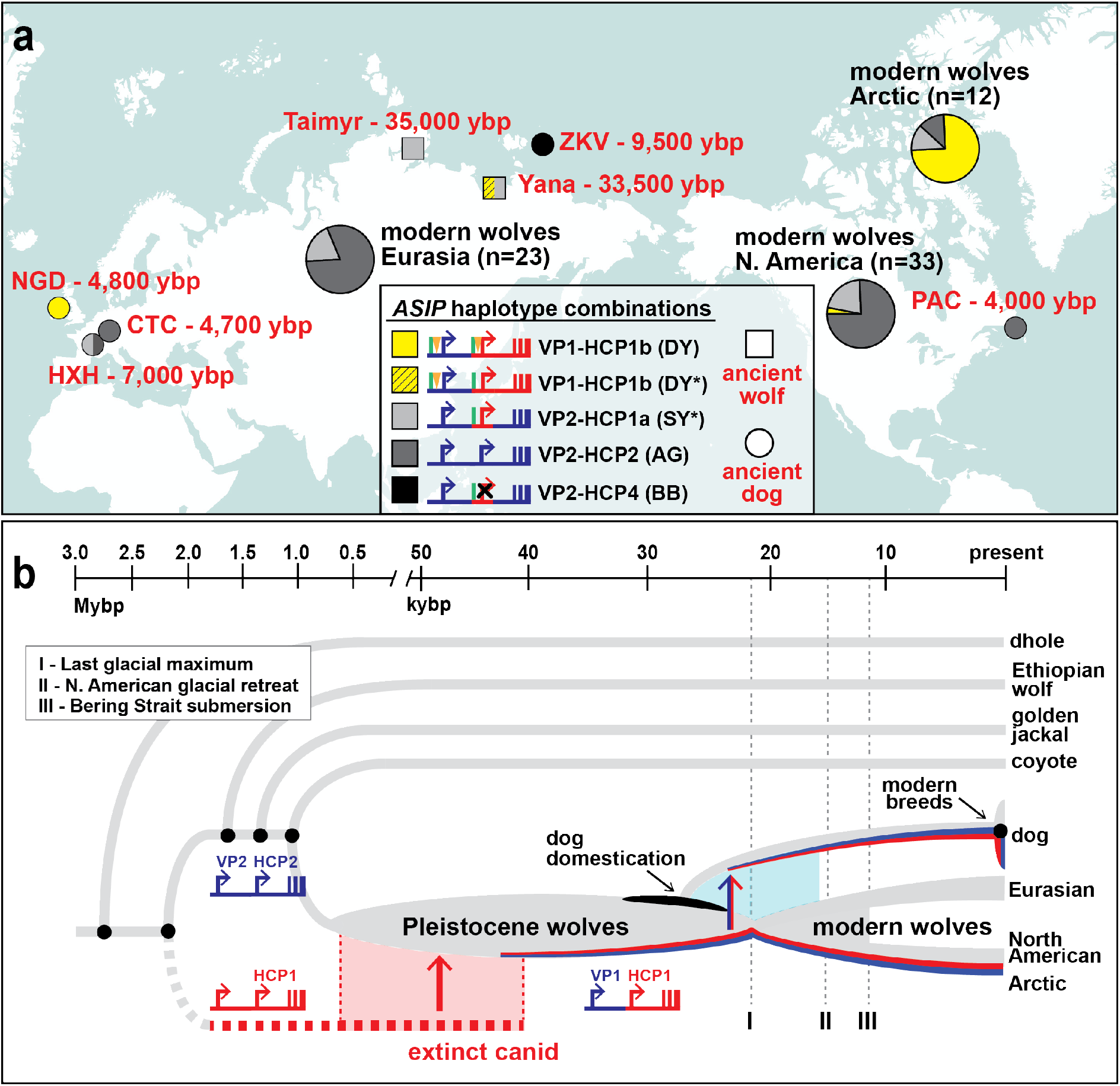
Distribution of *ASIP* alleles in ancient dogs and wolves, and an evolutionary model for dominant yellow acquisition. (**a**) *ASIP* haplotypes were inferred from whole genome sequencing of 5 ancient dog (circles), 2 ancient wolves (squares), and 68 modern wolves (pie charts) distributed across the Holarctic (see Supplementary Table 10 and Extended Data Fig. 8 for detailed haplotype representations). Asterisks indicate SY/DY haplotypes for which the HCP1 insertion is either absent (SY*) or not ascertainable (DY*). (**b**) A model for origin of the dominant yellow haplotype and its transmission into dogs and arctic wolves, in which molecular alterations at modular promoters were acquired by introgression (red, HCP1) or by mutation in the grey wolf (blue, VP1). The timeline for speciation events, dog domestication, and geological events affecting grey wolf dispersal are based on prior studies ^13,17^.

Together with our phylogenetic results, comparative analysis of wolf and dog *ASIP* haplotypes suggests an evolutionary history in which multiple derivative haplotypes and associated color patterns arose by recombination and mutation from two ancestral configurations corresponding to a white wolf (VP1-HCP1) and a grey wolf (VP2-HCP2), both present in the late Pleistocene (Fig. 4a, Extended Data Fig. 8). The distribution of derivative haplotypes explains color pattern diversity not only in dogs but also in modern wolf populations across the Holarctic, including white wolves in the North American Arctic (VP1-HCP1b) and yellow wolves in the Tibetan highlands (VP2-HCP1a), and is consistent with natural selection for light coat color.

A likely timeline for the origin of modules driving high levels of *ASIP* expression is depicted in Fig. 4b and indicates a dual origin. The HCP1 haplotype represents introgression into Pleistocene grey wolves from an extinct canid lineage that diverged from grey wolves more than 2 Mybp. This introgression as well as the mutation from VP2 to VP1 occurred prior to 33,500 ybp, based on direct observation from an ancient wolf sample (Fig. 4a). Natural selection for VP1 and HCP1 are a likely consequence of Pleistocene adaptation to arctic environments and genetic exchange in glacial refugia, driven by canid and megafaunal dispersal during interglacial periods. Modern grey wolves are thought to have arisen from a single source ∼25,000 ybp close to the last glacial maximum ^18,19^; during the North American glacial retreat that followed, the VP1-HCP1b haplotype combination was selected for in today’s white-colored arctic wolves.

In dogs, ASIP color pattern diversification was likely an early event during domestication, since our analysis of ancient DNA data reveals several different VP and HCP haplotypes in Eurasia by 4,800 ybp. This is consistent with the wide distribution of dominant yellow across modern dog breeds from diverse locations, as well as the dingo (Supplementary Table 9), a feral domesticate introduced to Australia at least 3,500 ybp ^20^. Of particular interest is the Zhokov island dog from Siberia ^21,22^. Based on a haplotype combination of VP2-HCP4, this sled dog that lived 9,500 years ago exhibited a black back color pattern, allowing it to be easily distinguished from white colored wolves in an arctic environment. Our results show how introgression, demographic history, and the genetic legacy of extinct canids played key roles in shaping this diversity.

## Methods

All data generated or analyzed during this study are included in this published article (and its supplementary information files).

### Ethics Statement

All animal experiments were done in accordance with the local regulations. Experiments were approved by the “Cantonal Committee For Animal Experiments” (Canton of Bern; permits 48/13, 75/16 and 71/19).

### Skin biopsies and total RNA extraction

Skin biopsies were taken from three dogs (a black back Miniature Pinscher and a dominant yellow Border Terrier and Irish Terrier). Two 6 mm punch biopsies were taken from differentially pigmented body areas of each animal (dorsal and ventral). RNA samples from dogs represent asynchronous hair growth relative to the hair cycle. The biopsies were immediately put in RNAlater (Qiagen) for at least 24 h and then frozen at –20°C. Prior to RNA extraction, the skin biopsies were homogenized mechanically with the TissueLyser II device from Qiagen. Total RNA was extracted from the homogenized tissue using the RNeasy Fibrous Tissue Mini Kit (Qiagen) according to the manufacturer’s instructions. RNA quality was assessed with a FragmentAnalyzer (Agilent) and the concentration was measured using a Qubit Fluorometer (ThermoFisher Scientific).

### Whole transcriptome sequencing (RNA-seq)

From each sample, 1 μg of high quality total RNA (RIN >9) was used for library preparation with the Illumina TruSeq Stranded mRNA kit. The libraries were pooled and sequenced on an S1 flow cell with 2×50 bp paired-end sequencing using an Illumina NovaSeq 6000 instrument. On average, 31.5 million paired-end reads per sample were collected. One publicly available Beagle sample was used (SRX1884098). All reads that passed quality control were mapped to the CanFam3.1 reference genome assembly using STAR aligner (version 2.6.0c) ^26^.

### Transcript coordinates

The STAR-aligned bam files were visualized in the IGV browser ^27^. Three different alternate untranslated first exons that appeared to splice to the coding exons of *ASIP* were defined based on the visualizations of the read alignments in IGV. These exact transcripts have not been documented in NCBI and Ensembl gene models. The visually curated gene models are given in Supplementary Table 2.

### Identification of genomic variants

WGS data from 71 dogs and 6 wolves was used for variant discovery (Supplementary Table 3). They included 15 agouti dogs and wolves, 25 black back dogs, 11 black saddle dogs, 14 dominant yellow dogs and 11 shaded yellow dogs and one white wolf. The genomes were either publicly available or sequenced as part of related projects in our group ^28^. SNVs and small indels were called as described ^28^. The IGV software ^27^ was used for visual inspection of the promoter regions based on the transcripts identified in the RNA sequencing data. Structural variants were identified and association with coat color phenotypes was verified by visual inspection in IGV.

### DNA samples for Sanger sequencing and genotyping

Samples for variant discovery included two dogs from each color phenotype and are designated in Supplementary Table 5 with asterisks. Samples from dogs listed in Supplementary Table 5 were used for genotyping. The coat color phenotype of all animals was assigned based on breed-specific coat color standards or photographs or owner reporting. Genomic DNA was isolated from EDTA blood samples using the Maxwell RSC Whole Blood DNA kit (Promega).

### Sequencing of promoter regions

Sanger sequencing of PCR amplicons was carried out to validate and characterize structural variants at the sequence level in the promoter regions. All primer sequences and polymerases used are listed in Supplementary Table 4. PCR products amplified using LA Taq polymerase (Takara) or Multiplex PCR Kit (Qiagen) were directly sequenced on an ABI 3730 capillary sequencer after treatment with exonuclease I and shrimp alkaline phosphatase. Sequence data were analyzed with Sequencher 5.1 (GeneCodes). Interspersed repeat insertions were classified with the RepeatMasker program ^29^. Multiple copies of SINE elements from the same and different families were resolved this way. The CanFam3.1 reference genome assembly is derived from the Boxer Tasha, a dominant yellow dog, and represents a DY haplotype, VP1-HCP1, of the *ASIP* gene. Descriptions of the promoter variants and Genbank accession numbers for HCP2-5 are in Supplementary Table 1. The table lists the 7 combinations of VP and HCP regulatory modules observed in dogs. As HCP3, HCP4, and HCP5 all represent loss-of-function alleles that are functionally equivalent, the 7 listed combinations correspond to only 5 distinct phenotypes.

### Genotyping assays

The previously reported SINE insertion ^24^ was genotyped by fragment size analysis on an ABI 3730 capillary sequencer and analyzed with the GeneMapper 4.0 software (Applied Biosystems). The previously reported *ASIP* coding variants ^25^ were genotyped by Sanger sequencing PCR products. The previously reported *RALY* intronic duplication ^23^ was genotyped by size differentiation of PCR products on a Fragment Analyzer (Agilent). Five PCR-assays (ventral promoter assays 1, 2; hair cycle promoter assays 1, 2, 3) are required to unambiguously determine the VP and HCP haplotypes. The other four primer pairs in the list were used to genotype previously published diagnostic markers ^23-25^ or for the amplification of the entire HCP (Supplementary Table 4). Genotyping results for all samples are shown in Supplementary Table 5. There is a perfect genotype-phenotype association in 352 dogs (see Fig. 2). In the remaining 14 dogs, the presence of a eumelanistic mask due to an epistatic *MC1R* allele prevented the reliable phenotypic differentiation of dominant yellow and shaded yellow dogs. Breeds and the different promoter haplotype combinations identified within each breed are indicated in Supplementary Table 6. In a few dogs that were heterozygous at both VP and HCP, the phasing of the VP and HCP haplotype combinations was performed based on haplotype frequency within the same breed as noted. A family of Chinooks were used to determine the segregation of extended haplotypes and the phenotypic equivalency of HCP3 and HCP5 (Extended Data Fig 3). Summary of genotyping results and exclusion of previously associated variants is shown in Supplementary Table 7. This table lists the genotype-phenotype association in aggregated form. The table also contains the genotypes for variants that were previously reported to be associated with pattern phenotypes ^23-25^. Numbers in red indicate genotyping results, for which these markers yielded discordant results.

### Comparison of promoter haplotype effects on transcripts

Transcript data was generated from a second set of samples. Sample descriptions and colors are shown in Supplementary Table 8 for all RNA experiments. Skin samples were collected from a male Swedish Elkhound (agouti), female German Pinscher (dominant yellow) and male Rottweiler (black back) after euthanasia that was conducted due to behavioral or health problems not related to skin. Samples were collected in RNAlater Stabilization Solution and stored at – 80°C. RNA was extracted using the RNeasy Fibrous Tissue Mini Kit (Qiagen) according to manufacturer’s instructions. Integrity of RNA was evaluated with Agilent 2100 Bioanalyzer or TapeStation system (Agilent) and concentration measured with DeNovix DS-11 Spectrophotometer (DeNovix Inc.). The libraries for STRT (Single cell reverse tagged) RNA-sequencing were prepared using STRT method with unique molecular identifiers ^30^ and modifications including longer UMI’s of 8 bp, addition of spike-in ERCC control RNA for normalization of expression, and Globin lock method ^31^ with LNA-primers for canine alpha- and betaglobin genes. The libraries were sequenced with an Illumina NextSeq 500. Reads were mapped to the CanFam3.1 genome build using HISAT1 mapper version 2.1.0 ^32^.

The alignment-free quantification method Kallisto (version 0.46.0) ^33^ was used to estimate the abundance and quantified as transcripts per million mapped reads (TPM) data based on an index built from CanFam3.1 Ensembl transcriptome (release 99). The curated *ASIP* transcript isoform models based upon alignment visualizations in the IGV browser ^27^ were also included in the transcriptome. Results based on genotype of the promoter haplotypes are displayed in Extended Data Fig. 1 as TPM.

### Haplotype construction

Haplotypes were constructed from two publicly available VCF files PRJEB32865 and PRJNA448733. The VCFs for selected dogs were merged using BCFtools merge tool (http://samtools.github.io/bcftools/) with the parameter --missing-to-ref, which assumed genotypes at missing sites are homozygous reference type 0/0. Only dogs homozygous for ASIP haplotypes were used to visualize haplotypes (Supplementary Table 3). SNVs that had 100% call rate in these samples were color coded and displayed relative to the genome assembly and previously commercialized variants (Extended Data Fig. 2).

### *ASIP* phylogenetic analysis in canids

Illumina whole genome sequence for 36 canids, including seven extant species and the dog, were downloaded from the NCBI short read archive as aligned (bam format) or unaligned (fastq format reads (Supplementary Table 9). Fastq data were aligned to the dog genome (CanFam3.1) using BWA (v.0.7.17) ^34^ after trimming with Trim Galore (v.0.6.4). SNVs within a 110 kb interval (chr24:23,300,000-23,410,000), which includes the *ASIP* transcriptional unit and regulatory sequences, were identified with Platypus (v.0.8.1) ^35^ and filtered with VCFtools (v.0.1.15) ^36^ to include 2008 biallelic SNVs. Phasing was inferred with BEAGLE (v.4.1) ^37^.

For phylogenetic analysis, the *ASIP* interval was partitioned in two regions, based on dog SNV density (Fig. 3a) and *ASIP* gene structure: a 48 kb region including the ventral first exon, extending to but excluding the hair cycle first exon (chr24:23,330,000-23,378,000), and a 16 kb region including the hair cycle first exon, extending to and including *ASIP* coding exons 2-4 (chr24:23,378,001-23,394,000). Consensus sequences of equal length were constructed for each inferred canid haplotype using BCFtools (v.1.9). Phylogenies were inferred using Maximum Likelihood method and Tamura-Nei model with 250 bootstrap replications, implemented in MEGAX ^38,39^, and including 34 canids (Fig. 3b, Extended Data Fig. 4,5). For 34 of 36 individuals, consensus haplotype pairs were adjacent to each other or, in the case of a few wolf/dog haplotypes, were positioned in neighboring branches with weak bootstrap support. The exceptions were the African golden wolf, a species derived by recent hybridization of the grey wolf and Ethiopian wolf ^12^, and an eastern grey wolf from the Great Lakes region, which was also reported to have recent admixture with the coyote ^40^. The African golden wolf and the eastern grey wolf were removed from the alignments, and a single haplotype for each individual was selected arbitrarily for tree building and display.

### Haplotype analysis of *ASIP* locus in ancient dogs and wolves

Whole genome sequencing data from several recent studies ^12,16,22,41-45^, including five ancient dogs, two ancient grey wolves, and 68 modern grey wolves (Supplementary Table 10) were downloaded as aligned (bam format) or unaligned (fastq format) reads. Fastq data was aligned to the dog genome (canFam3.1) using BWA-MEM (v.0.7.17) ^34^, after trimming withTrim Galore (v.0.6.4). Coverage depth for each sample ranged from 1-78x (Supplementary Table 10). Genotypes at five structural variants and six SNVs were determined by visual inspection using the IGV browser (Supplementary Table 10). Variants in or near the ventral promoter (n=2), the hair cycle promoter (n=6), and the coding exons (n=3) distinguished ventral and hair cycle promoter haplotypes (Supplementary Table 10, Extended Data Fig. 7). SNV genotypes were determined by allele counts; structural variants were genotyped by split reads at breakpoint junctions.

For 67 of 75 wolves (or ancient dogs), the phase of ventral and hair cycle promoter haplotypes was unambiguous. Seven wolves and one ancient dog were heterozygous with respect to both the ventral and hair cycle promoter haplotypes, and for these samples, haplotype phase was inferred based on the linkage disequilibrium in the 67 unambiguous individuals.

## Acknowledgements

We would like to acknowledge the Next Generation Sequencing Platform of the University of Bern and Biomedicum Functional Genomics Unit (FuGU), University of Helsinki, for sequencing services and the Interfaculty Bioinformatics Unit of the University of Bern and IT Center For Science Ltd. (CSC, Finland) for providing high performance computing infrastructure. We thank resources and members of the Dog Genome Annotation (DoGA) Consortium (Hannes Lohi, Juha Kere, Carsten Daub, Marjo Hytönen, César L. Araujo, Ileana B. Quintero, Kaisa Kyöstilä, Maria Kaukonen, Meharji Arumilli, Milla Salonen, Riika Sarviaho, Julia Niskanen, Sruthi Hundi, Jenni Puurunen, Sini Sulkama, Sini Karjalainen, Antti Sukura, Pernilla Syrjä, Niina Airas, Henna Pekkarinen, Ilona Kareinen, Anna Knuuttila, Heli Nordgren, Karoliina Hagner, Tarja Pääkkönen, Kaarel Krjutskov, Sini Ezer, Shintaro Katayama, Masahito Yoshihara, Auli Saarinen, Abdul Kadir Mukarram, Matthias Hörtenhuber, Amitha Raman, Irene Stevens) as well as the Dog Biomedical Variant Database Consortium and all other canine researchers who deposited genome sequencing data into public databases. We thank the dog owners who provided photographs.

## Author Contributions

DB: conceptualization, investigation, writing, visualization, formal analysis, CK: investigation, visualization, formal analysis, writing, AL, PH, RL: validation, resources, VJ and MR: software, PR, JH: validation, KM and JM: resources, MKH, AM, HL, DoGA consortium: resources, STRT analyses, CD: supervision and resources, GB: supervision, writing-review and editing, TL: conceptualization, funding acquisition, investigation, supervision, resources, writing-review and editing.

## Additional Information

Supplementary Information is available for this paper

Correspondence and requests for materials should be addressed to Danika L. Bannasch

Reprints and permissions information is available at www.nature.com/reprints

## Extended data

**Extended Data Fig. 1:**
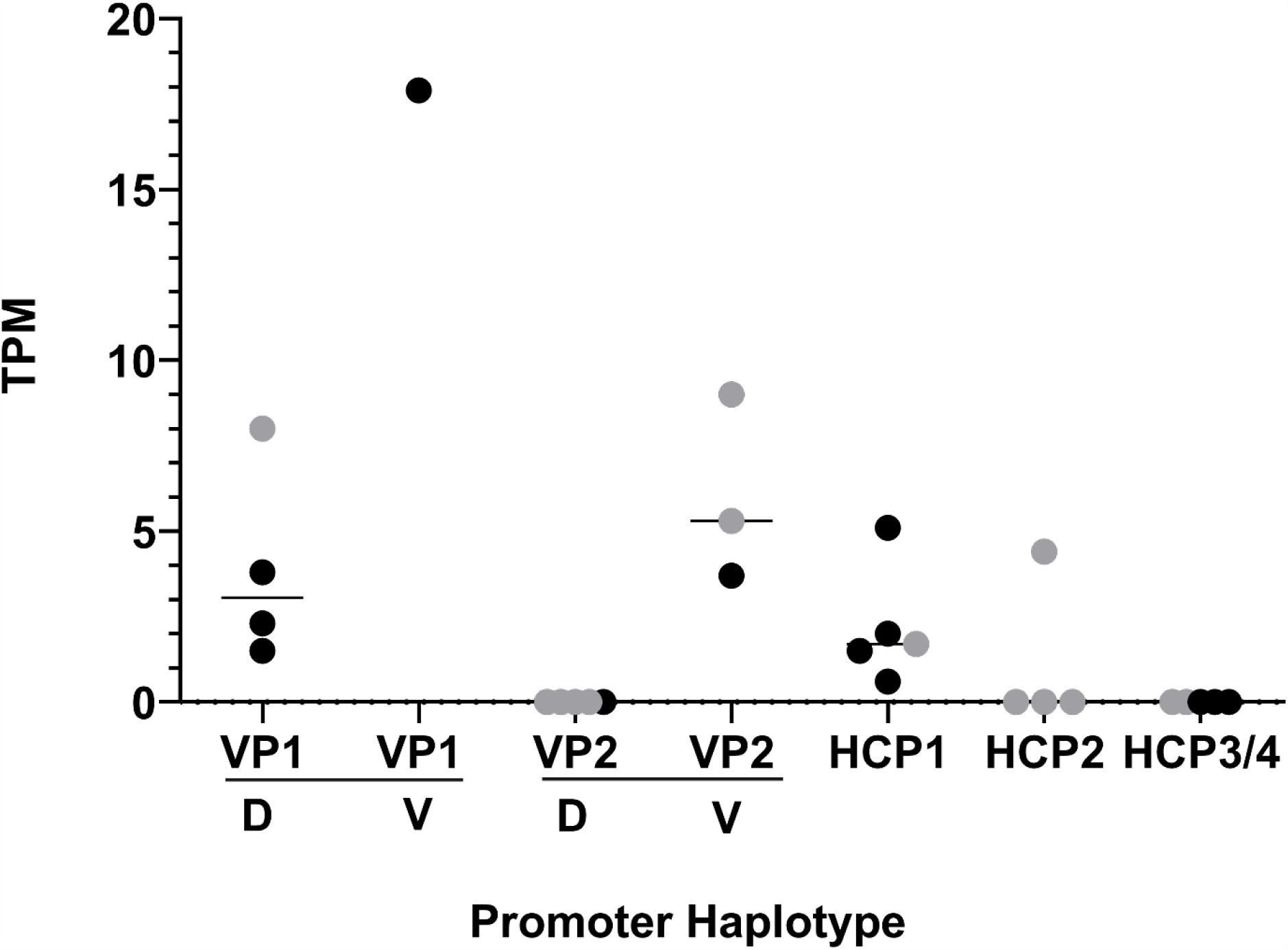
Relative transcription of promoter variants. Black dots are from RNA-seq data and grey dots are from STRT RNA-seq data.Dorsal samples (D) were taken from mid thorax of the dog and ventral (V) from the ventral abdomen. The HCP samples were not synchronized with respect to the hair cycle.

**Extended Data Fig. 2:**
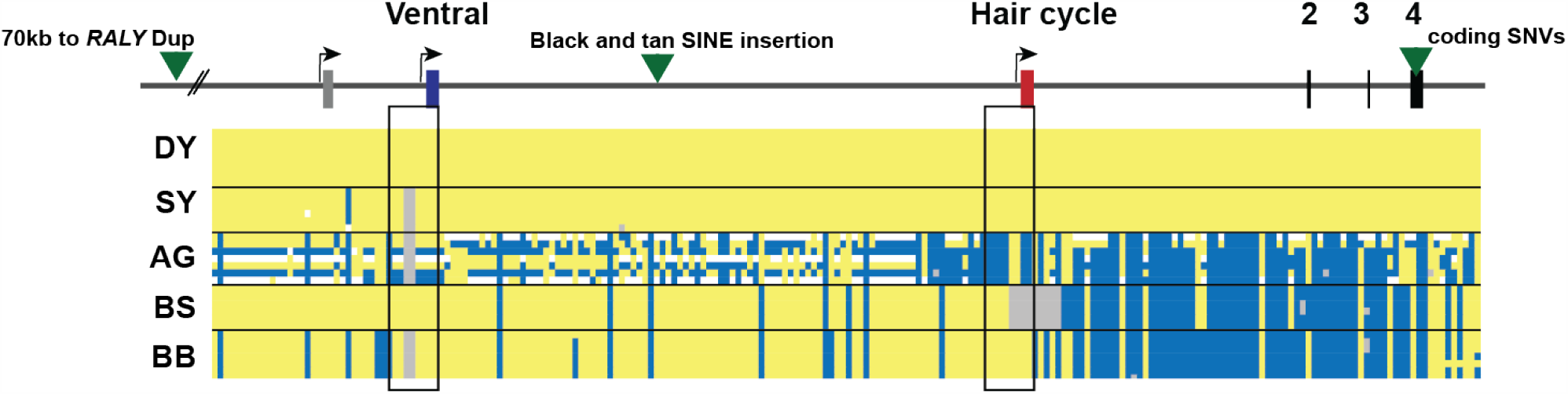
Dog haplotypes across the *ASIP* locus with comparison to commercial genetic tests for coat color. Dog coat pattern phenotypes are listed on the left. Alternative first exons are listed at the top. Yellow is a homozygous match to the genome assembly, grey heterozygous, white deleted and blue homozygous alternate allele. The black rectangles highlight the promoter regions. Green triangles represent the location of variants that were previously used in commercial testing to distinguish different alleles for coat color patterns. The previously identified intronic duplication that was promoted to commercially distinguish BS and BB haplotypes in some breeds lies 70 kb to the left of this diagram ^23^. The green triangle between the VP and HCP is the location of the commercially tested SINE insertion for BB and BS ^24^. In the samples presented here, the dominant yellow haplotype extends through the coding sequence where the missense variants associated with this haplotype were previously identified ^25^. In more primitive breeds, recombination events have disrupted this long linkage disequilibrium between the promoter variants and the coding variants leading to incorrect genetic test results with the existing tests. Samples used are listed in Supplementary Table 3. Raw genotyping results are in Supplementary Table 5 and summary results comparing commercial variants are in Supplementary Table 7.

**Extended Data Fig. 3:**
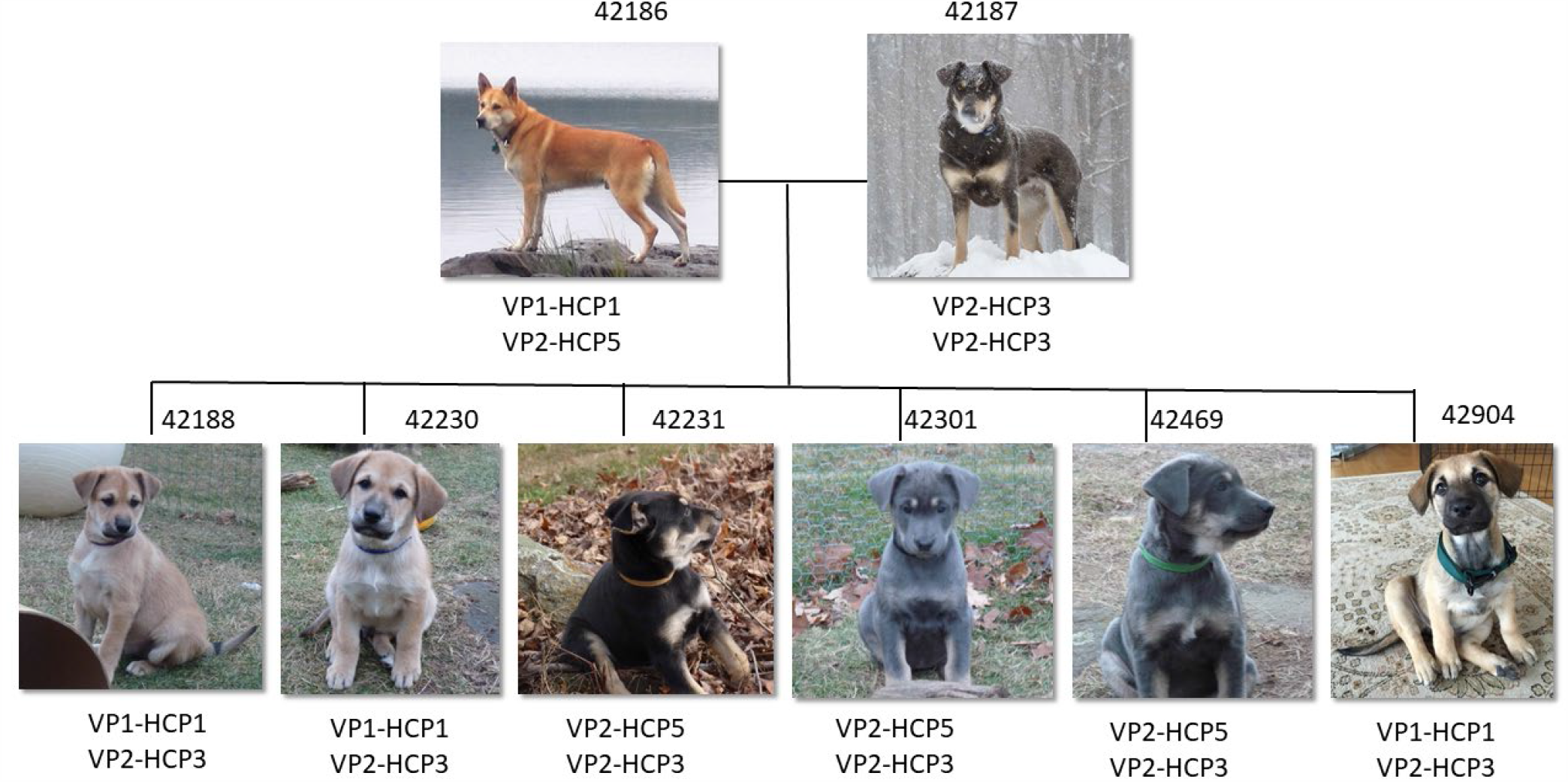
A family of dogs segregating dominant yellow and black back. Extended haplotype combinations were determined in this family of Chinook dogs. In this breed both HCP3 and HCP5 segregate and confer a black back phenotype combined with VP2.

**Extended Data Fig. 4:**
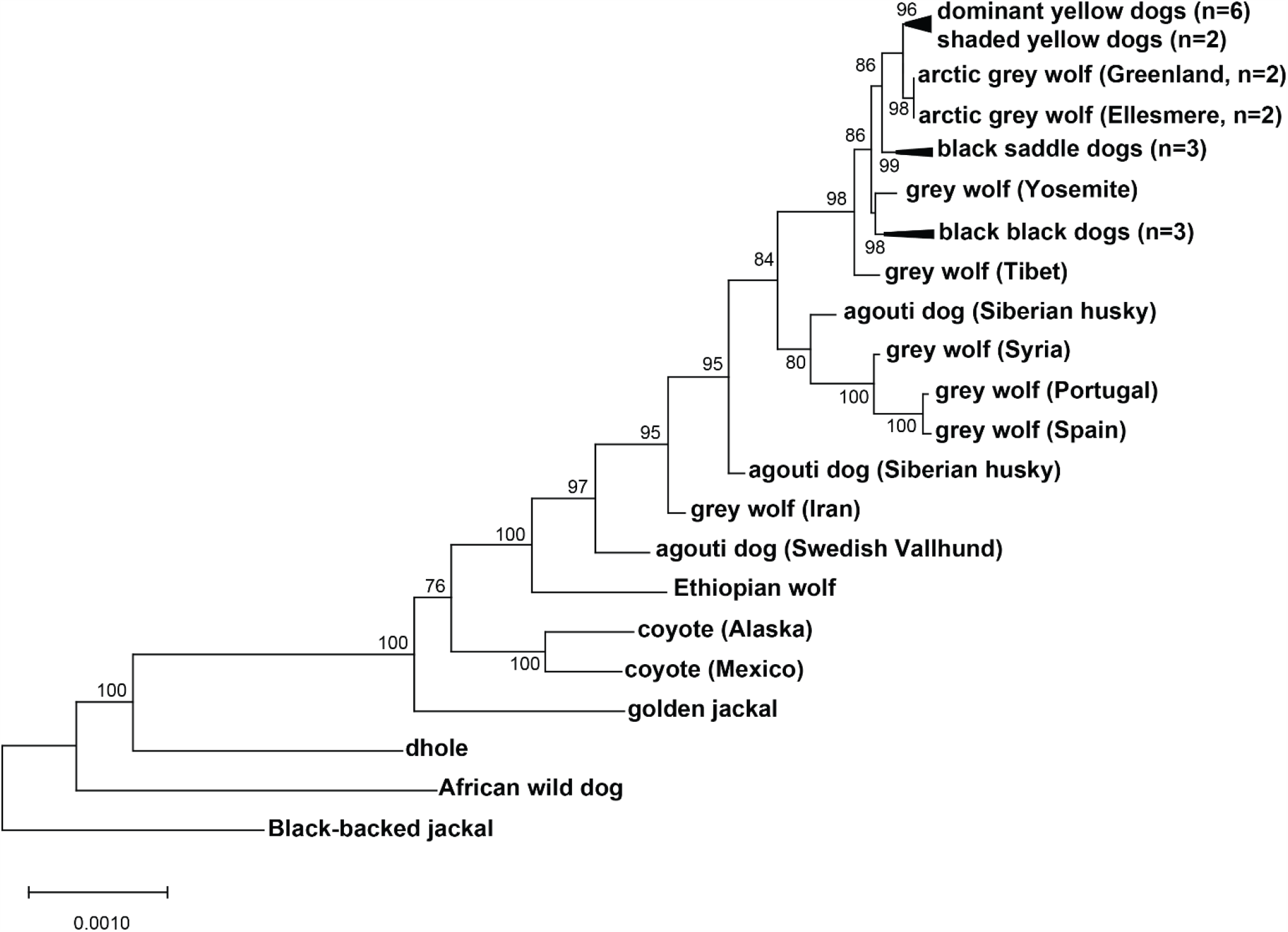
Expanded canid phylogenetic tree inferred from 48 kb region including the ventral promoter. An expanded version of the maximum likelihood tree shown in Fig. 3B, with 34 canids, representing 7 of 9 extant species.

**Extended Data Fig. 5:**
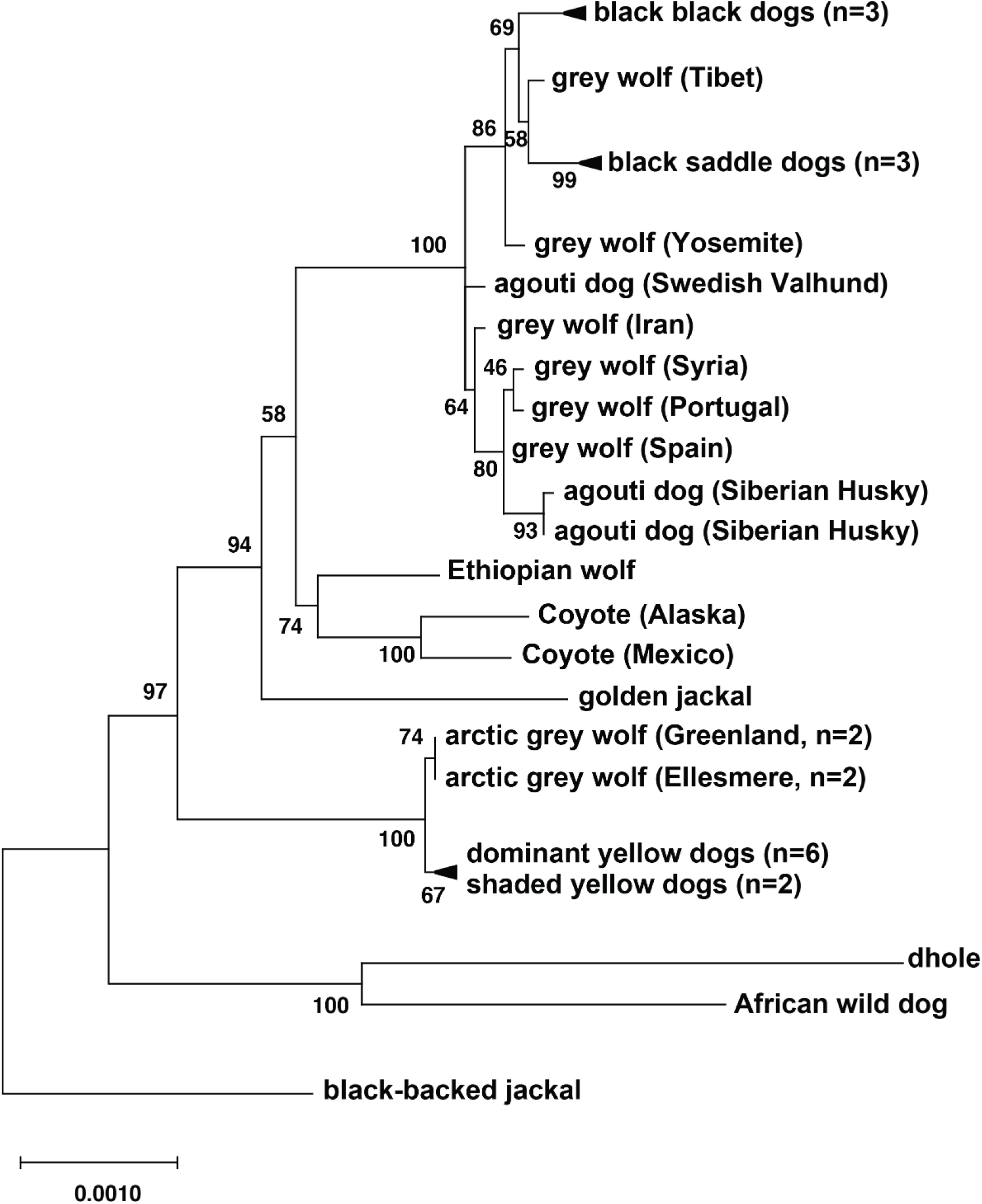
Expanded canid phylogenetic tree inferred from 16 kb region within and downstream of the hair cycle promoter. An expanded version of the maximum likelihood tree shown in Fig. 3b, with 34 canids, representing 7 of 9 extant species.

**Extended Data Fig. 6:**
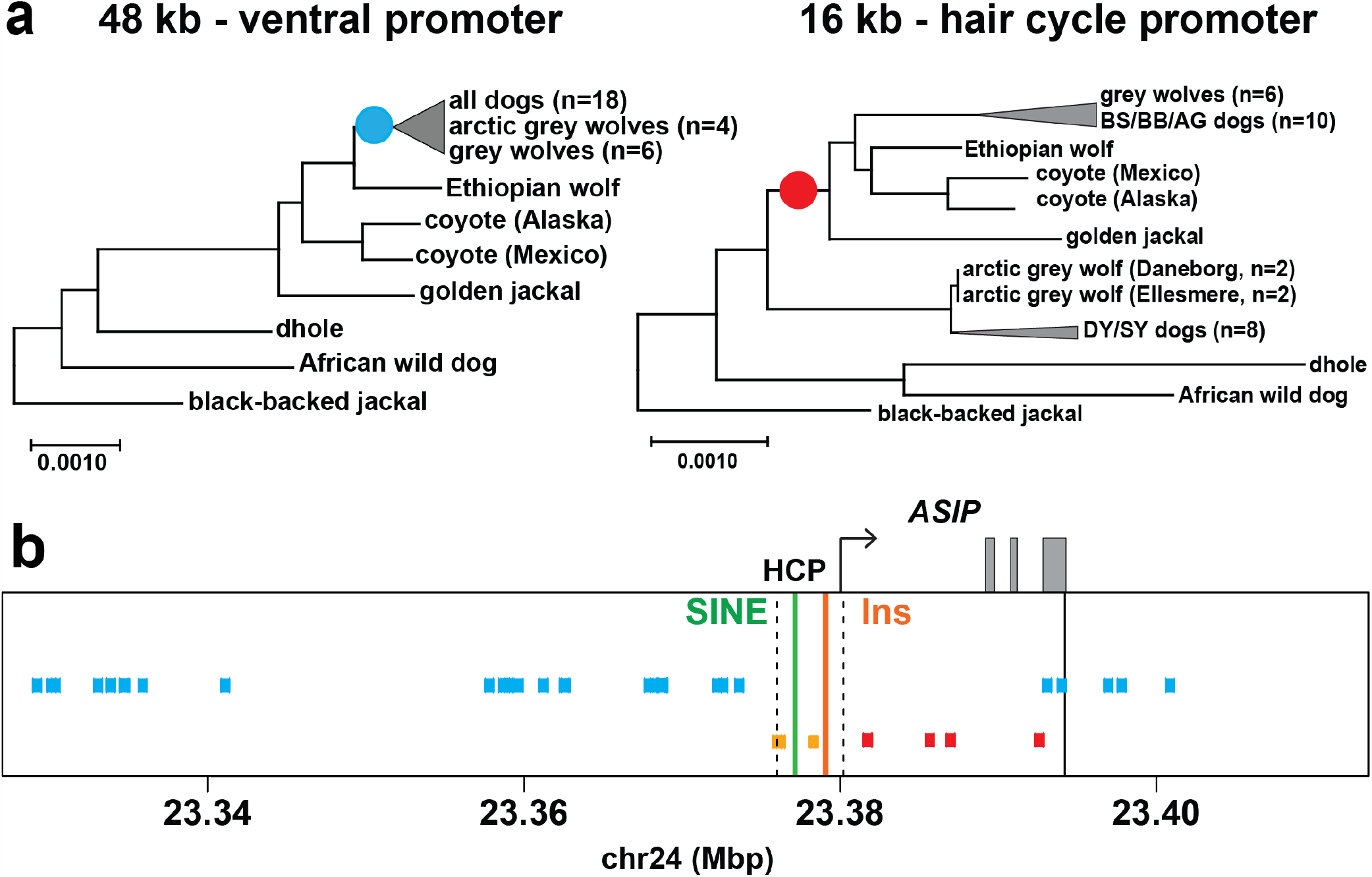
Genomic distribution of derived substitutions across the *ASIP* locus. (**a**) Canid phylogenies for the ventral (48 kb) and hair cycle (16 kb) promoter regions, with relevant internal branches marked by the occurrence of derived variants plotted in (B). (**b**) Derived substitutions shared by grey wolf and dogs (cyan). Ancestral alleles on DY/arctic wolf haplotypes (red) or BB and DY/arctic wolf haplotypes (orange) that correspond to derived substitutions in the core wolf-like canids (Supplementary Table 11). The broken lines demarcate the HCP region (chr24:23,375,800-23,380,000). The solid line signifies the downstream boundary for phylogenetic analysis. The solid green and orange lines indicate the positions of the SINE and 24 bp insertion, respectively, associated with the DY/arctic wolf haplotype.

**Extended Data Fig. 7:**
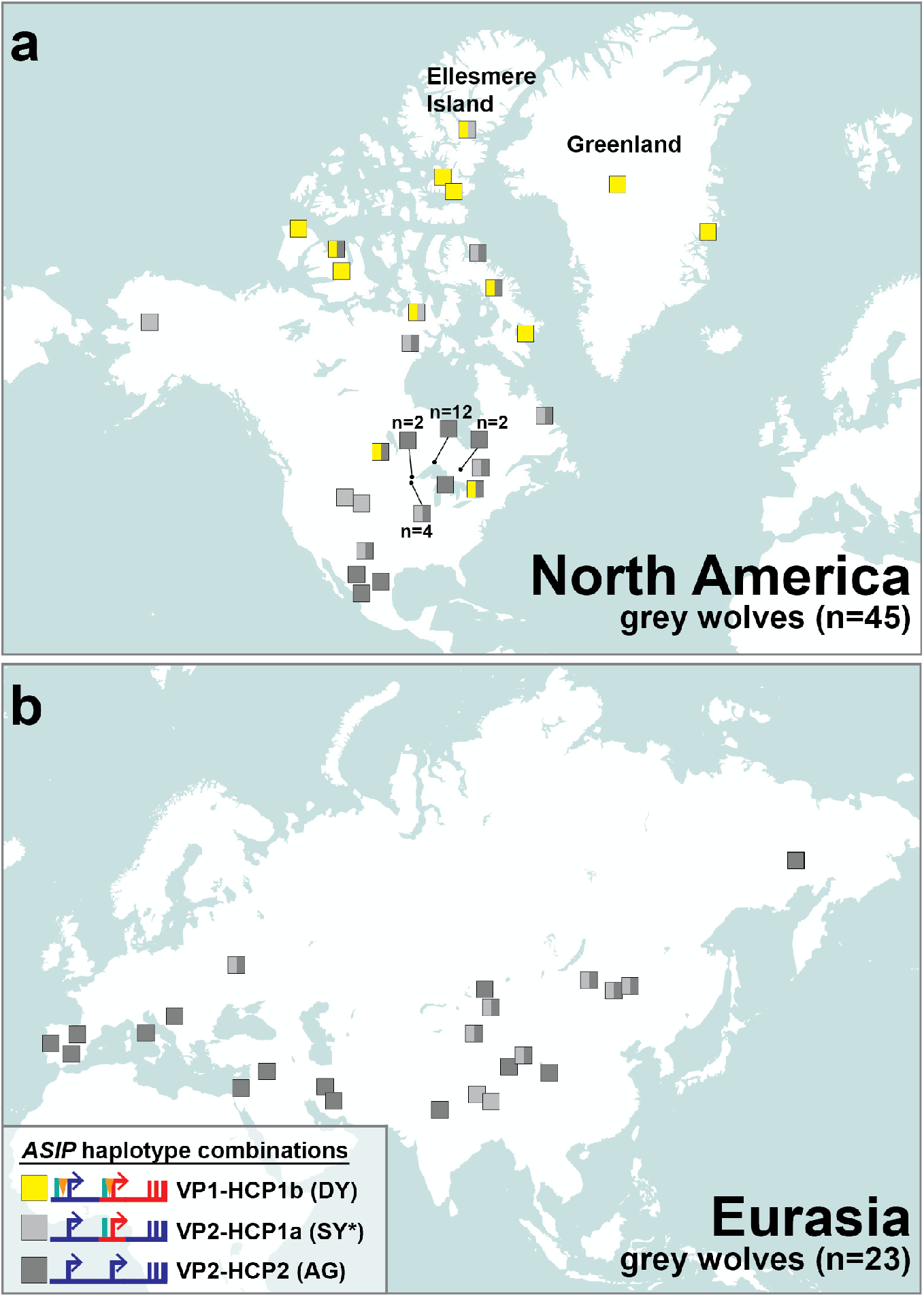
The distribution of *ASIP* haplotypes in modern grey wolves. Modern grey wolves from (a) North America (n=45) or (b) Eurasia (n=23) were genotyped for 5 structural variants and 6 SNVs using whole genome sequencing data. Each wolf, represented by a colored box, is plotted information, summarized in the figure legend, is available in Extended Data Fig. 8 and Supplementary Table 10. The asterisk indicates an SY-like haplotype without the HCP1 insertion.

**Extended Data Fig. 8:**
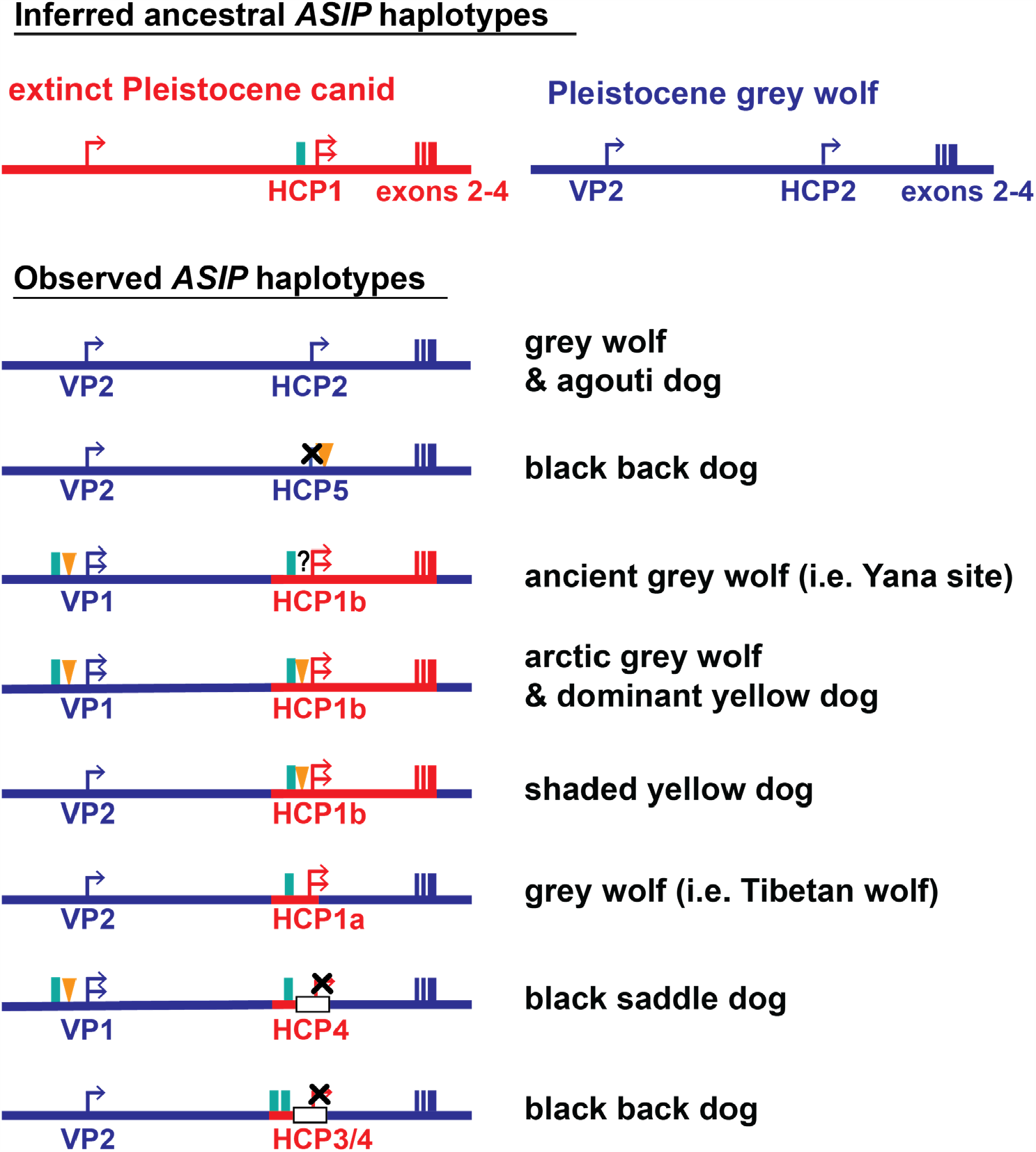
Evolutionary diversification of *ASIP* haplotypes observed in grey wolves and dogs. The color (red or blue) of *ASIP* haplotype segments indicates ancestral species of origin, inferred from phylogenetic analysis (Fig. 3b, Extended Data Fig. 4, 5). Relevant structural variants near the ventral (VP) and hair cycle (HCP) promoters are depicted as yellow triangles (polynucleotide expansions), green bars (SINE insertions), and white bars (deletions). Modified promoter activity is indicated by an X mark (no activity) or an additional arrow (elevated expression), based on RNAseq (Extended Data Fig. 1) and/or inference from coat color (Fig. 1, 3c).

**Extended Data Table 1.** Segregation of modular promoter diplotypes with phenotype.

| Phenotype | Diplotype | Counts |
| --- | --- | --- |
| Dominant Yellow | VP1-HCP1 / VP1,2-HCP1,3,4,5<br>VP2-HCP1 / VP2-HCP1 | 113/114<br>1*/114 |
| Shaded Yellow | VP2-HCP1 / VP2-HCP1,3,5<br>VP2-HCP1 / VP1-HCP1<br>VP1-HCP1 / VP1-HCP1 | 51/64<br>11*/64<br>2*/64 |
| Agouti | VP2-HCP2 / VP2-HCP2,3,5 | 46/46 |
| Black Saddle | VP1-HCP4 / VP1,2-HCP4,3 | 53/53 |
| Black Back | VP2-HCP3 / VP2-HCP3,4,5 | 89/89 |

\* Dogs had MC1R based eumelanin masking pattern, which prevented reliable phenotype distinction between dominant yellow and shaded yellow.

**Extended Data Table 2.** SNVs distinguishing DY dogs and arctic wolves in the 64kb segment that contain the VP, HCP, and coding sequences.

| Position<br>(Chr24, CF3) | Ancestral<br>allele | Derived<br>Allele | DY dog | Arctic wolf | Yana wolf |
| --- | --- | --- | --- | --- | --- |
| 23,333,763 | C | A | C | A | C |
| 23,343,447 | G | C | C | G | G |
| 23,356,213 | T | C | T | C | T |
| 23,362,891 | C | A | C | A | C |
| 23,381,935 | C | T | C | T | C |
| 23,393,514 | G | A | A | G | G |

