## Supplemental Information for "Dog color patterns explained by modular promoters of ancient canid origin"

**Supplementary Information**

**Supplementary Tables 1-11**

**Text**

***ASIP* locus phylogenetic analysis**

Maximum likelihood trees constructed from ventral (48 kb) and hair cycle (16 kb) promoter regions were consistent with genome-wide analysis ^1^, with three exceptions (Fig. 3b). Two involved incongruities apparent in both trees. First, the African golden wolf, a species derived from admixture of grey and Ethiopian wolves ^1^, had distinct *ASIP* haplotypes clustering with or between its ancestral species and was excluded from phylogenies. Second, the Ethiopian wolf, basal to the golden jackal in genome-wide phylogenies ^1,2^, is inferred to be a sister group to either the grey wolf (ventral promoter region) or coyote (hair cycle promoter region), possibly revealing ancient admixture. The third exception, as described in the main text, is apparent only in the hair cycle promoter tree, as predicted by the skewed distribution of sequence variation among dog and wolf haplotypes (Fig. 3a). In the ventral promoter tree, all dogs (n=18) and grey wolves (n=10) form a single clade with strong bootstrap support. However, in the hair cycle promoter tree, arctic grey wolves (n=4) and DY/SY dogs (n=8) form a separate clade that is basal to the golden jackal and distinct from other canid species.

**Evolutionary history of the dominant yellow haplotype**

Characterization of *ASIP* structural variation within promoter regions of 8 canid species indicates that the agouti dog haplotype (VP2-HCP2) is ancestral within the canid lineage (Fig. 3d). VP2 and HCP2 structure is shared by all non-dog canids, with the exception of some grey wolves, discussed in detail below.

The most prominent exceptions are the arctic grey wolves from Ellesmere Island and Greenland (n =4), all of which share extended haplotypes with dominant yellow dogs (VP1-HCP1), spanning the entire transcriptional unit and including all derived structural variants at the ventral and hair cycle promoters. All haplotypes from these four high arctic wolves are identical except for one polymorphic site, and are distinguished from dog dominant yellow haplotypes by 6 SNVs across 64 kb (Extended Data Table 2). Notably, the arctic wolf haplotype has only one of the two derived *ASIP* coding variants (NC_006606.3:g.23393514A>G) present on the dog dominant yellow haplotype. The haplotype architecture of dominant yellow dogs and arctic grey wolves suggests a common origin without recent genetic exchange.

A closely related haplotype (DY*, Fig. 4a and Extended Data Fig. 7) was also present in an ancient wolf sample, dating to 33.5 kybp, from the Yana River in Northern Siberia. The inferred DY* is nearly identical to haplotypes in dominant yellow dogs and arctic wolves, including at least three of the four derived SINE and polynucleotide insertions at *ASIP* promoters (Extended Data Fig. 8, Supplementary Table 10). No reads span the HCP 24 bp insertion. Only ancestral alleles were observed at each of the 6 SNV loci that distinguish the dominant yellow and arctic wolf haplotypes (Extended Data Table 2), indicating that high latitude Pleistocene wolves harbored a haplotype ancestral to those present in modern yellow dogs and white wolves.

The first evidence of the dominant yellow haplotype in a dog is from an ancient DNA sample from the Newgrange tomb in Donore, Ireland, dated to 4900-4700 ybp (Fig. 4a). The Newgrange dog was homozygous for the same DY haplotype present in contemporary dogs, indicating that yellow coat color in dogs traces to at least the Neolithic period, consistent with other evidence including the estimated arrival of the dingo, a predominantly yellow, feral domesticate in Australia, at least between 3363-3211 ybp^3^. This evidence provides a lower bound for the presence of the dominant yellow haplotype in dogs, but given its inferred dispersal across Eurasia in the Neolithic, its introduction into dogs likely occurred much earlier, perhaps during domestication.

**Extinct canid introgression haplotypes at the *ASIP* locus**

To refine the extinct canid introgression haplotype, we examined the pattern of derived allele sharing among wolf-like canids. Derived alleles shared by all dogs and grey wolves but not other canids, i.e. those occurring along the phyletic branch indicated in the left panel of Extended Data Fig. 6a (cyan), mark haplotype segments corresponding to a contemporary wolf lineage. Alternatively, derived alleles shared by wolf-like canids (grey wolf, coyote, golden jackal, and Ethiopian wolf) but not arctic wolves and DY/SY dogs, i.e. those occurring along the phyletic branch indicated in the right panel of Extended Data Fig. 6a (red), reveal haplotype segments corresponding to an introgression. We identified 37 and 9 SNVs corresponding to the former and latter phyletic relationships, respectively (Extended Data Fig. 6b, Supplementary Table 11). The distribution of these derived alleles across the *ASIP* locus delineates a minimal introgression haplotype that, in arctic wolves and DY/SY dogs, extends 16.5 kb (chr24:23,375,968-23,392,505, CanFam3.1).

Four of the nine SNVs that delineate the introgression haplotype are located within the hair cycle promoter region (orange circles in Extended Data Fig. 7a, chr24:23,375,968-23,378,833), and are in perfect linkage disequilibrium with a DY-associated HCP SINE insertion (Fig. 3d, Extended Data Fig. 7, Supplementary Table 10). Together these variants can be used to detect recombinant introgression haplotypes which do not extend distally to *ASIP* coding exons. HCP3 and HCP4, which harbor the promoter impairing deletions responsible for black back, occur on the introgression haplotype (Fig. 2, 3d, and Extended Data Fig. 8). The observation of the HCP4 haplotype in an ancient dog, dating to 9,500 ybp ^4^ excavated from Zhokhov Island in the East Siberia Sea, indicates existence of the introgression haplotype and black back coat pattern in neolithic dogs. Archeological evidence suggests Zhokhov Island dogs were selectively bred for body size variation ^5^. The observed HCP4 haplotype suggests coat color was also a target for selection.

We also identified a recombinant introgression haplotype in contemporary wolves, including 7 of 8 Tibetan wolves, a distinct light-colored grey wolf subspecies (*Canis lupus chanco*) adapted to high altitude Tibetan and Himalayan plains (Fig. 3d and Extended Data Fig. 7). The Tibetan wolf HCP1 haplotype lacks the 24 bp polynucleotide expansion present in dominant yellow dogs and arctic grey wolves; we refer to these two closely related HCP1 haplotypes as HCP1a and HCP1b, respectively (Fig. 3d and Extended Data Fig. 8).

Characterization of HCP structural variants present on introgression haplotypes in modern wolves and dogs reveals the chronological order of variant acquisition and the evolutionary relationship of HCP haplotypes (Fig. 3d). The SINE insertion shared by HCP1, 3, and 4 likely predates introgression into Pleistocene wolves. An additional SINE insertion occurred along the lineage that gave rise to HCP3 and HCP4, prior to the deletions that impaired promoter activity. The 24 bp polynucleotide expansion is present only on HCP1b and therefore occurred after the SINE insertion but before transmission to the dog and the arctic wolf (Fig. 3d).

**Likelihood Dominant Yellow haplotype acquisition by incomplete lineage sorting**

Nine derived SNVs located within and downstream of the hair cycle promoter region are shared by four wolf-like canid species but not DY/SY dogs and arctic wolves (Extended Data Fig. 6). The pattern of allele sharing strongly supports an introgression from an extinct canid lineage, but does not explicitly exclude the possibility that the haplotype arose in a direct grey wolf ancestor and persisted through multiple speciation events as a consequence of incomplete lineage sorting and balancing selection. In the latter scenario, the DY haplotype would be subject to gradual decay by recombination. To test the likelihood of DY haplotype acquisition by ancestral lineage transmission, we applied a probability distribution for haplotype decay previously used to test incomplete lineage sorting in other contexts ^6^. Specifically, we determined the expectation that a 16.5 kb haplotype could persist in the grey wolf lineage for the length of time (*t*) that predates the haplotype delineating sequence variation, given the average grey wolf genome-wide recombination rate. We applied the lower bound of the estimated split time between the Ethiopian and grey wolf (1.5 Mybp, ^2^) as a conservative estimate of haplotype divergence time, a recombination rate (*r*) of 0.78 cM/Mb ^7^, and generation time of 3 years. The expected mean length (*L*) of a persisting haplotype is 1/(*r* × *t*), and the probability of observing a haplotype ≥ length *m* fits a Gamma distribution with shape parameter 2, and rate parameter 1/*L* ^6^. The expected size of a canid haplotype persisting since the split of the Ethiopian and grey wolf is 256 bp, and the likelihood of observing a 16.5 kb haplotype is *p* = 0. Notably, a scenario in which introgression occurs during the late Pleistocene (i.e. ~50 kybp) is consistent with a Gamma distribution model of decay by recombination (*p*=0.37).
